# Chemotherapy Induces an IL1β-dependent Neutrophil Recruitment that Promotes Chemoresistance in Metastatic Ovarian Cancer

**DOI:** 10.1101/2025.09.21.677600

**Authors:** Taito Miyamoto, Yujie Ye, Bryan Manning, Brennah Murphy, Moeko Minakuchi, Kohei Hamada, Rin Mizuno, Mana Taki, Koji Yamanoi, Ryusuke Murakami, Masaki Mandai, Ying Ye, Jayamanna Wickramasinghe, Yulia Nefedova, Andrew Kossenkov, Nan Zhang

**Author notes:** Correspondence to: Nan Zhang and Taito Miyamoto, Nan Zhang, PhD Assistant Professor, Molecular and Cellular Oncogenesis Program Ellen and Ronald Caplan Cancer Center, The Wistar Institute, Taito Miyamoto, MD, PhD Assistant Professor, Department of Gynecology and Obstetrics Kyoto University Graduate School of Medicine. Equal contribution (Taito Miyamoto, Yujie Ye). **Conflict of interest disclosure statement** The authors declare that there is no conflict of interest regarding the publication of this article.

## Abstract

High-grade serous carcinoma (HGSC) of the ovary acquires chemoresistance through diverse cancer cell-intrinsic and-extrinsic mechanisms, culminating in treatment-refractory intraperitoneal metastasis. How chemotherapy-induced remodeling of the tumor microenvironment modulates drug sensitivity remains unclear. In this study, we demonstrate that chemotherapy induced IL1β-dependent neutrophil accumulation in tumors, driving chemoresistance in HGSC. Using patient samples, bulk transcriptomic profiling before and after chemotherapy revealed post-treatment upregulation of *IL1B*, and single-cell RNA sequencing identified myeloid cells as its principal source. In a chemoresistant murine metastatic ovarian cancer model, chemotherapy increased neutrophils and neutrophil extracellular traps (NETs) in omentum tumors; these increases were abrogated in IL1β-deficient mice, with expansion of activated CD8+ T cells and tumor control. Neutrophil depletion in wild-type mice recapitulated the chemosensitive phenotype of IL1β-deficient mice. *In vitro*, IL1β did not alter cancer cell-intrinsic chemosensitivity, whereas NETs reduced the chemosensitivity of cancer cells. Additionally, the dominant IL1β receptor (IL1R1) was predominantly expressed in tumor-associated fibroblasts in humans and mice. Consistently, IL1R1-deficient mice exhibited chemosensitivity with decreased neutrophil accumulation and increased IFNγ⁺TNF⁺CD8⁺ T cells. We also found that chemotherapy upregulated CXCL2 in patients and that ablating IL1β-IL1R1 axis decreased CXCL2 expression in tumor-associated fibroblasts in mice. Finally, residual human HGSC tumor after chemotherapy showed increased neutrophils and a trend toward more NETs. Collectively, these findings illuminate a paradoxical, cancer cell-extrinsic mechanism in HGSC whereby chemotherapy itself amplifies chemoresistance and suggest that targeting chemotherapy-induced inflammation may help overcome treatment resistance.

## Introduction

Ovarian cancer (OC) causes more than 200,000 deaths annually worldwide and ranks as the eighth leading cause of cancer-related mortality among women (1). OC is commonly diagnosed at an advanced stage with peritoneal dissemination, necessitating chemotherapy-based multimodal treatment including bevacizumab and PARP inhibitors (2,3). Nevertheless, the majority of advanced-stage patients ultimately experience chemotherapy-resistant tumor regrowth (4,5). Overcoming chemoresistance therefore remains a critical challenge in the OC management.

As the most prevalent subtype of OC, high-grade serous carcinoma (HGSC) exhibits extensive genomic instability and copy number alterations, primarily associated with TP53 mutations (6). Cancer cell intrinsic mechanisms of chemoresistance—namely tumor–specific genomic and epigenomic alterations and transcriptional programs—have been recognized as a key contributor to treatment failure. Primary resistance has been associated with copy number amplifications of *CCNE1* and *KRAS* (7,8). Moreover, therapy-induced acquired resistance involves reversion mutations in *BRCA1/2* (7,9) and demethylation of the *BRCA1* promoter (7,10). However, these alterations are observed only in a subset of patients. Moreover, reports indicating minimal genomic differences between primary and recurrent tumors (8) suggest that chemoresistance in HGSC is not solely determined by specific genomic or epigenomic profiles of cancer cells, but rather represents a multifactorial phenotype shaped by additional contributory mechanisms.

In contrast, cancer cell-extrinsic mechanisms of chemoresistance—namely, those arising from indirect modulation of cancer cells or the formation of physical barriers by components of the tumor microenvironment (TME), such as vasculature, immune cells, and fibroblasts—have been proposed (11–13). In HGSC, primary chemoresistance has been associated with factors such as increased stromal content and the presence of regulatory T cells (14,15). Importantly, the TME is not static but undergoes dynamic remodeling during chemotherapy; however, the extent to which chemotherapy-induced changes in the TME influence treatment responses remains poorly understood.

In this study, we demonstrate that, in the context of chemotherapy, myeloid cell–derived IL1β drives neutrophil recruitment, thereby fostering chemoresistance in HGSC. This observation reveals a paradoxical mechanism whereby chemotherapy-induced inflammatory remodeling of the TME impairs tumor sensitivity to chemotherapy. Our study sheds light on one aspect of the multifactorial nature of chemoresistance in HGSC and suggests the potential of overcoming treatment resistance through inhibition of therapy-induced inflammation.

## Materials and Methods

### Bulk RNA sequencing analysis of human OC samples before-and after-chemotherapy

Gene expression data from pre-and post-chemotherapy ovarian cancer tissue samples derived from the same patients were obtained from the following datasets: Javellana et al., 2022 (16), GSE109934 (17), GSE181597 (18), and GSE201600 (19). Among these, the dataset from Javellana et al. consists of RNA-seq data, while the remaining three are based on targeted gene expression panels. For the RNA-seq dataset from Javellana et al., paired differential expression was performed with limma R package (RRID: SCR_001905) (20) on log2-transformed values, and genes meeting FDR < 0.05 and ≥1.5-fold change were analyzed in IPA (QIAGEN; Upstream Regulators). Inflammasome activity was quantified by ssGSEA (GenePattern v3.5.0; RRID: SCR_003201) with MSigDB REACTOME_INFLAMMASOMES and THE_NLRP3_INFLAMMASOME gene sets following the methodology described by Barbie et al (21). We further analyzed gene expressions of *IL1B* and *TNF* in pre-and post-chemotherapy samples from GSE109934, GSE181597, and GSE201600. In GSE181597, *IL1B* expression was additionally stratified and compared by chemotherapy response score (CRS) classification. *CXCL1*, *CXCL2*, *CXCL3*, *CXCL5*, *CXCL6*, *CXCL8*, *CSF2*, and *CSF3*—cytokines implicated in neutrophil migration and differentiation—were assessed in relevant datasets, except GSE109934, which did not contain chemokine expression data.

### Single cell RNA sequencing analysis of human OC samples before-and after-chemotherapy

The treatment-naïve scRNA-seq dataset generated by Vázquez-García et al. was retrieved from the Gene Expression Omnibus (GSE180661) (22). A scRNA-seq dataset from post-chemotherapy patients was retrieved from the study by Zhang et al. (GSE165897) (23). The Seurat package v4.3.0 (RRID:SCR_016341) in R software v4.3.1 and the scanpy package v1.9.8 (RRID:SCR_018139) in Python 3.10 (RRID:SCR_008394) were used for downstream processing. Dimensionality reduction was performed using principal component analysis and uniform manifold approximation and projection (UMAP) following the original protocol, with cell-type annotations from the original article. The R package scCustomize v1.1.1 (RRID:SCR_024675) was used to generate density and joint plots.

### KM Plotter

The KM Plotter Online tool (https://kmplot.com/analysis/) (24) was used to evaluate the relationship between gene expression and clinical outcome in patients with OC. This open-access TCGA-based database contains bulk RNA sequencing datasets from 374 OC patients, which allowed us to investigate correlation between overall survival (OS) or progression free survival (PFS) and *IL1B*.

### Cell lines

KPCA and BPCA cells, which were recently generated and characterized (Iyer et al., 2021) (25), were generously gifted to us by Dr. Robert Weinberg at the Whitehead Institute. All cell lines were cultured in Dulbecco’s modified Eagle’s medium (DMEM, 10-017-CV, Corning) supplemented with 4% FBS (#16140-071, Gibco), 1% penicillin/streptomycin (#15140122, Gibco), 1X Insulin-Transferrin-Selenium (ITS, #41400-045, Gibco), and 2 ng/mL mouse epidermal growth factor (mEGF, #315-09, Gibco) at 37°C supplied with 5% CO_2_. Cells were passaged no more than 15 times prior to injection into mouse peritoneal cavities.

### Animal models

C57BL/6J (WT, RRID:IMSR_JAX:000664), C57BL/6J-*Il1b^em2Lutzy^*/Mmjax (IL1β KO, RRID:MMRRC_068082-JAX), and B6.129S7-*Il1r1^tm1Imx^*/J (IL1R1 KO, RRID:IMSR_JAX:003245) mice were purchased from Jackson Laboratories or MMRRC. Mice were kept on a 12hr light-dark cycle and had access to food and water *ad libitum*. All animal procedures were performed in accordance with the Wistar Institutional Animal Care and Use Committee under protocol 201536.

### Tumor implantation, treatment, and evaluation

Cells were harvested with trypsin-EDTA (25-052-CI, Corning), washed in PBS, and injected intraperitoneally (i.p.) into mice. For KPCA tumors, 1×10^6^ KPCA cells were injected i.p. into WT, IL1β KO, or IL1R1 KO mice in 200μL of a 1:1 matrigel:PBS mix (Matrigel Matrix Basement Membrane, #354234, Corning). At day 7, mice were treated with 10mg/kg carboplatin (C2538, Sigma) and 1mg/kg paclitaxel (NSC 125973, Selleck) or vehicle control once a week for two weeks. At day 21, mice were imaged by IVIS and applied for necropsy analysis next day. For neutrophil depletion studies, the mixture of 200μg of anti-Ly6G (L280, Leinco Tech, RRID:AB_2737551) and 50μg of anti-rat kappa light chain (I-2027, Leinco Tech,) monoclonal antibodies (26) were injected i.p. in WT or IL1β KO KPCA tumor-bearing mice together with chemotherapy at day 7 and then 3 times a week for 2 weeks. For BPCA tumors, 3×10^6^ BPCA cells were injected i.p. into WT or IL1β KO mice in 200μL of a 1:1 matrigel:PBS mix. Mice were treated with 10mg/kg carboplatin and 1mg/kg paclitaxel or vehicle control at day 35 and day 49. At day 64, mice were applied for necropsy analysis.

### *In vivo* Bioluminescent Imaging

*In vivo* bioluminescence imaging was performed on an IVIS 50 (PerkinElmer; Living Image 4.3.1), with exposures of 1 s to 1 min, binning 2–8, field of view 12.5 cm, f/stop 1, and open filter. D-Luciferin (150 mg/kg in PBS, eLUCNA-1G, Gold Biotechnology) was injected into the mice i.p. and imaged 10min later.

### Mesentery Metastasis Scoring

Mesentery metastasis score was calculated as previously reported (27). Briefly, all mesenteric tumor nodules were counted and each mouse score was determined as following:

0: no tumor was detected,

1: number of tumor nodules is less than 10, 2: number of nodules is 10-30,

3: number of nodules is over 30.

### Tissue dissociation

Omentum tumors were digested using a cocktail of 1mg/ml collagenase IV (C5138, Sigma) and 100 µg/ml DNase I (NC1839861, Sigma), in 2-3mL RPMI with 10% FBS. The tissues were minced into small pieces and digested at 37°C for 30 min shaking at 800 rpm with intermittent vortexing. Samples were then passed through a 70µm cell strainer to collect single-cell suspension to be analyzed by flow cytometry.

### Flow cytometry

Single-cell suspensions were collected as described above prior to staining with primary conjugated antibodies at their indicated dilutions (Supplementary Table 1). Intracellular staining was carried out using True-Nuclear™ Transcription Factor Buffer Set (#424401, Biolegend). Mesenchymal populations were defined within CD45^-^ non-immune populations as follows: fibroblast (Podoplanin^+^PDGFRα^+^CD31^-^), mesothelial cells (Podoplanin^+^PDGFRα^-^CD31^-^) and endothelial cells (Podoplanin^-^PDGFRα^-^CD31^+^). For IFNγ and TNF staining, single cells were incubated for 4 hour with Cell Activation Cocktail (with Brefeldin A) (#423303, Biolegend) prior to staining. Samples were analyzed on a BD FACSymphony™ A3 Cell Analyzer using FlowJo software (RRID:SCR_008520).

### Cell viability assay with chemotherapeutic agents in the presence or absence of IL1β

On Day 0, KPCA cells were seeded at a density of 2,000 cells per well in 96-well clear flat-bottom plate. On Day 1, the culture medium was replaced with fresh medium containing either carboplatin at final concentrations of 0.1, 10, 50, 100, 200, or 500 μM, or paclitaxel at 0.01, 0.1, 1, 2, 10, or 1000 nM. In each condition, recombinant mouse IL1β (401-ML-005, R & D) was co-administered at concentrations of 0, 0.1, 1, or 10 ng/mL. On Day 4, cell viability was assessed using CCK-8 solution (HY-K0301, MedChemExpress).

### Single cell sequencing and analysis of mouse tumors

From omental tumors (n=3 mice/group), non-B cells were sorted and pooled. Samples of each group were then loaded onto a 10x Genomics Chromium Next GEM Single Cell 3′ v3.1 cartridge to generate droplets. Libraries were prepared per 10x instructions, QC’d (Bioanalyzer 2100, High Sensitivity DNA) and quantified (KAPA qPCR), then sequenced on an Illumina NextSeq 2000 (P3 100-cycle; paired-end: R1 28 bp, i7 8 bp, R2 90 bp) (RRID:SCR_023614).

BCL files were converted to FASTQ with Cell Ranger (v9.0.0, RRID:SCR_017344). Reads were aligned via STAR (v2.7.9a, RRID:SCR_004463) to the mm10 reference (10x refdata-gex-mm10-2020-A) and UMI counts were generated. Count matrices were imported into R and processed with Seurat (v5.3.0). Data were normalized, scaled, and analyzed on HVGs by PCA (top 50 PCs), embedded by UMAP, and clustered with the Louvain algorithm. Cell types were annotated with SingleR (RRID:SCR_023120) using ImmGenData (2024-02-26) and celldex (v1.18.0). Differential expression used Seurat’s Wilcoxon rank-sum test with Bonferroni correction. Visualizations (UMAPs, heatmaps) were generated in R (ggplot2).

### Immunofluorescence

Details of the antibodies used and their concentrations are provided in Supplementary Table S1. Mouse omentum was fixed with 4% paraformaldehyde. The 10μm-thickness tissue slides were blocked with 3% BSA and 1% Triton in PBS, and incubated with MPO and cHH3 primary antibodies overnight. The tissues were washed and incubated with respective secondary antibodies. After washing, incubate specimen in 300 nM DAPI solution. Confocal images were collected using a Leica SP8 microscope (RRID:SCR_018169).

Three-micrometer-thick formalin-fixed, paraffin-embedded (FFPE) sections of human omentum were stained using a sequential double staining protocol. After deparaffinization/rehydration, sections underwent sequential tyramide signal amplification (TSA)-immunofluorescence. MPO was labeled with primary antibody, HRP-polymer secondary, and TSA-Fluorescein Green (SAT701001EA, Akoya Biosciences); residual antibodies/HRP were inactivated by pH-9 antigen retrieval (#415201, Nichirei Biosciences). cHH3 was then detected using the same scheme with TSA-Cy5 Red (SAT705A001EA, Akoya Biosciences). DAPI was used for nuclear counterstain.

### Preparation of NETs-conditioned media

Bone marrow cells were isolated from the femurs and tibiae of 6–10-week-old female C57BL/6 mice. Neutrophils were enriched using a biotin positive selection kit (#17683, Stemcell Technologies) with biotin-conjugated Ly6G antibody (#127603, Biolegend, RRID:AB_1186105). Isolated neutrophils were resuspended in DMEM with 4% FBS at a concentration of 2 × 10⁵ cells/well (in 500 µL per well) and plated in 24-well plates. NET formation was induced by treating cells with 20 nM phorbol 12-myristate 13-acetate (PMA, P1585, Sigma). DMSO was used to make respective control media. In some conditions, the PAD4 inhibitor GSK484 (SML1658, Sigma) was added at a final concentration of 10 µM. After overnight incubation, supernatants (NETs-conditioned media) were collected by centrifugation.

### Cell viability assay with chemotherapeutic agents in NETs-conditioned media

KPCA cells were seeded at 2,000 cells/well in a 96-well clear flat-bottom plate. Next day, media was replaced with 100 µL of conditioned media from neutrophil cultures. Chemotherapeutic agents were added to designated wells: carboplatin (final concentration 20 µM) and paclitaxel (final concentration 10 nM), respectively. Cells were incubated for 48 hours at 37°C with 5% CO₂. After 48 hours, 30 µg of D-Luciferin was added to each well. Luminescence imaging and quantification were performed to assess cell viability.

### Tumor Immune Estimation Resource (TIMER)

We assessed the association between *IL1B*, *IL1R1* and *CXCL2* mRNA expression and immune infiltration using the TIMER 2.0 web platform http://timer.cistrome.org/ (28). Neutrophil abundance was inferred with the TIMER deconvolution algorithm on TCGA data. Spearman correlation coefficients between each gene expression and inferred neutrophil levels was computed adjusting for tumor purity.

### Human samples

Paired FFPE omental tumor samples from 16 HGSC patients at Kyoto University were collected before and after chemotherapy. Pre-treatment specimens were obtained at diagnostic laparotomy/laparoscopy; post-treatment specimens were collected at interval debulking surgery after carboplatin–taxane chemotherapy. From each block, 3-µm sections were prepared; adjacent sections underwent H&E and immunofluorescence staining. Neutrophils and NETs were quantified as described below. Peripheral-blood neutrophil counts were also compared between baseline (before chemotherapy) and immediately before interval debulking surgery (after chemotherapy). This analysis was approved by the Kyoto University Graduate School and Faculty of Medicine Ethics Committee (reference number G531) and conforms to the Declaration of Helsinki. Informed consent was obtained from all participants via an opt-in approach or an opt-out approach.

### Neutrophil quantification using HoVer-Next

For nuclear identification and cell type classification on H&E-stained slides, we employed HoVer-NeXt, a state-of-the-art AI model (29). HoVer-NeXt was trained on a modified version of the Lizard dataset and performs seven-class classification of the following cell types: (1) epithelial cells, (2) connective tissue cells, (3) lymphocytes, (4) plasma cells, (5) neutrophils, (6) eosinophils, and (7) mitotic figures. The whole-slide image was used as the region of interest, and default settings and parameters were applied for analysis. In the non-cancerous state, epithelial cells are absent in the omentum; therefore, epithelial cells identified within omental tumors are considered synonymous with cancer cells.

### NETs quantification using QuPath

For quantitative analysis of fluorescently labeled cells using multiplex immunofluorescence, QuPath (v0.6.0, RRID:SCR_018257) was utilized (30). The whole-slide image was used as the region of interest. Cell counts for DAPI, FITC (MPO), and Cy5 (cHH3) channels were obtained using the Cell Detection module, with default settings and thresholds applied throughout the analysis. NETs were defined by the colocalization of MPO and cHH3 (31). For each slide, the ratio of NETs to DAPI-positive cells was quantified.

### Statistics

Statistical analyses were performed in Prism (GraphPad Software, Inc., RRID:SCR_002798). The specific statistical tests applied—student *t* test, paired *t* test, or one-way ANOVA followed by Tukey’s post hoc multiple comparisons—are detailed in the figure legends. For mouse experiments, mice were randomly allocated to control and experimental groups. The investigator was not blinded to the group allocation. *A priori* power analysis was not performed. Results were considered significant at P < 0.05. Results are presented as mean ± SEM.

## Data availability statement

The mouse sequencing data is being submitted to the Gene Expression Omnibus. Remaining data are available from the corresponding author upon reasonable request.

## Results

### IL1β pathway is upregulated in post-chemotherapy human OC tumors

To investigate how chemotherapy alters the TME, we analyzed publicly available bulk RNA-sequencing datasets from patient-matched pre-and post-chemotherapy HGSC samples (16) and performed Ingenuity Pathway Analysis (IPA) with the focus on cytokine regulators. It revealed activation of inflammatory pathways, including TNF, IL1β, and IL6, as top hits in post-chemotherapy tumor samples (Figure 1A). Consistently, single-sample gene set enrichment analysis (ssGSEA) demonstrated upregulation of the inflammasome pathway, a key executioner for IL1β secretion, following chemotherapy (Figure 1B, 1C). Using other independent datasets, we next compared the gene expression levels of *TNF* and *IL1B*. Although *TNF*, the top upregulated cytokine in the IPA, showed no significant change between pre-and post-chemotherapy samples (Supplementary Figure S1A–1C), *IL1B*, the second most upregulated cytokine, exhibited increased expression following chemotherapy (Figure 1D, 1E). Furthermore, in the GSE181597 dataset, stratification by chemotherapy response revealed that *IL1B* expression remained unchanged in chemotherapy-sensitive cases (CRS3) but was notably elevated after treatment in chemotherapy-resistant cases (CRS1 and CRS2) (Supplementary Figure S1D–1F). To identify the cellular source of IL1β in HGSC, we analyzed single-cell RNA sequencing (scRNAseq) data from untreated and post-chemotherapy tumor samples (GSE180661 and GSE165897, respectively). Interestingly, *IL1B* expression was predominantly observed in myeloid cells, regardless of chemotherapy exposure (red arrows in Figure 1F and 1G). Finally, survival analysis of the TCGA-OV cohort using KM Plotter demonstrated that high *IL1B* expression in pretreatment tumor samples was associated with shorter overall survival (OS) and progression-free survival (PFS) among chemotherapy-treated HGSC patients (Figure 1H). Collectively, these findings suggest that chemotherapy-induced upregulation of *IL1B* may play a role in diminishing treatment sensitivity in HGSC.

**Figure 1.**
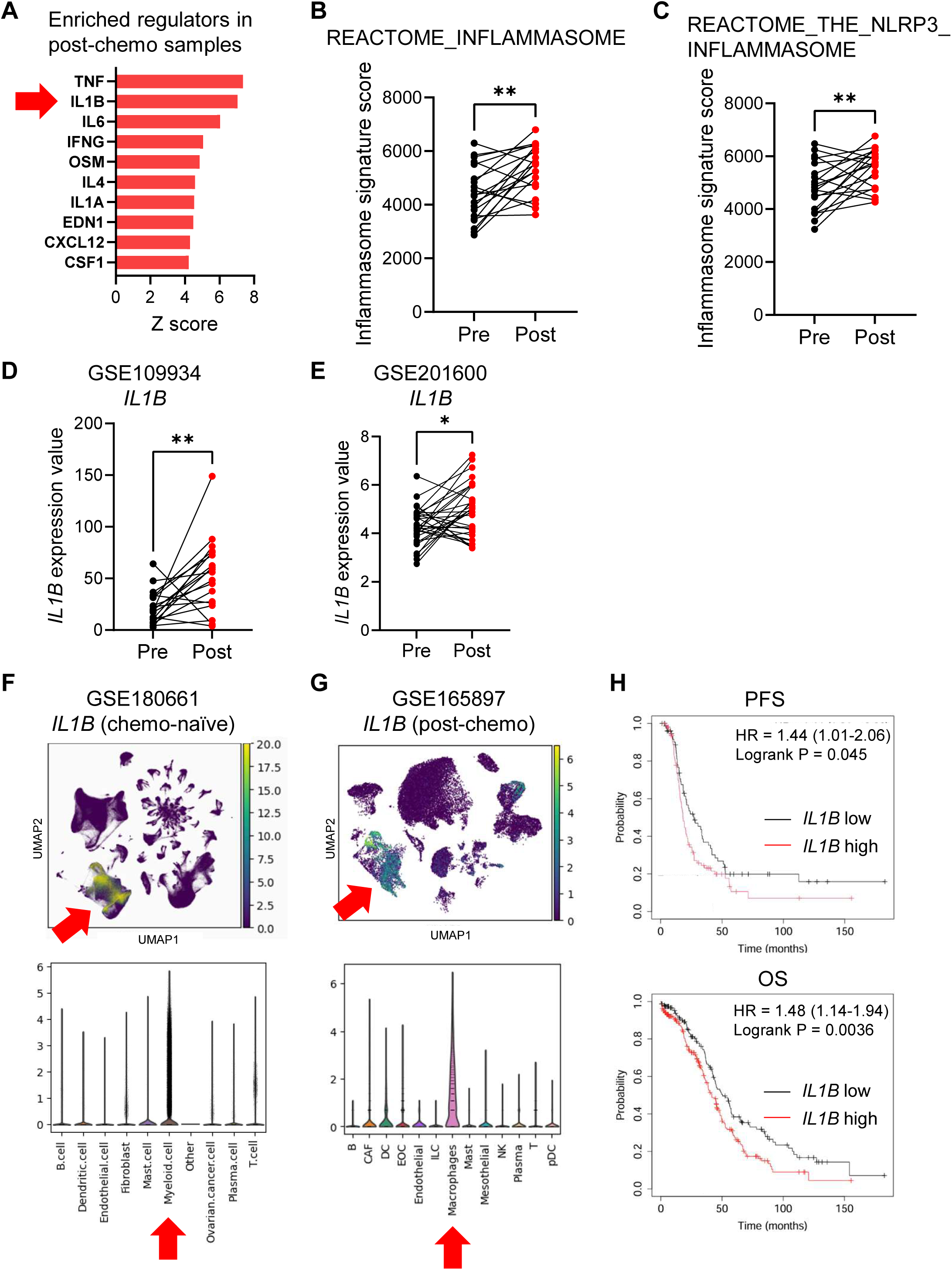
IL1β pathway is upregulated in post-chemotherapy human OC tumors. (A) Top 10 significantly enriched (FDR<5%) cytokine regulators predicted to be activated in post-chemotherapy samples by IPA. Red arrow highlights IL1β. (B and C) Comparison of inflammasome pathway signature scores between pre-and post-chemotherapy samples using (B) REACTOME_INFLAMMASOME gene set and (C) REACTOME_THE_NLRP3_INFLAMMASOME gene set. (D and E) Comparison of *IL1B* gene expression between pre-and post-chemotherapy samples using (D) GSE109934 dataset and (E) GSE201600 dataset. (F and G) Feature plot for *IL1B* analyzing (F) GSE180661 (dataset from treatment-naïve samples) and (G) GSE165897 (post-chemotherapy samples). Red arrows highlight myeloid cell populations. (H) Progression free survival (PFS) and overall survival (OS) analysis in OC patients with high and low expression of *IL1B*. Paired *t* test was used. *P < 0.05; **P < 0.01.

### IL1β from myeloid cells contributes to OC chemoresistance

We next sought to validate these findings using a murine OC model. To this end, we employed KPCA, a homologous recombination (HR)-proficient OC cell line derived from mouse fallopian tube epithelial cells that harbors genetic alterations relevant to human HGSC, including *CCNE1* amplification (25). An intraperitoneal dissemination model was established using KPCA cells, followed by chemotherapy with carboplatin and paclitaxel (Figure 2A). KPCA tumors exhibited resistance to chemotherapy (Figure 2B). As we previously reported, the HR-deficient cell line BPPNM demonstrated sensitivity to chemotherapy under similar conditions (27). Flow cytometry and scRNAseq analyses of IL1β expression across cell populations within omental tumors revealed that, consistent with the aforementioned human data (Figure 1F and 1G), IL1β was predominantly expressed by myeloid cells, i.e., monocytes, macrophages, and neutrophils (Figure 2C).

**Figure 2.**
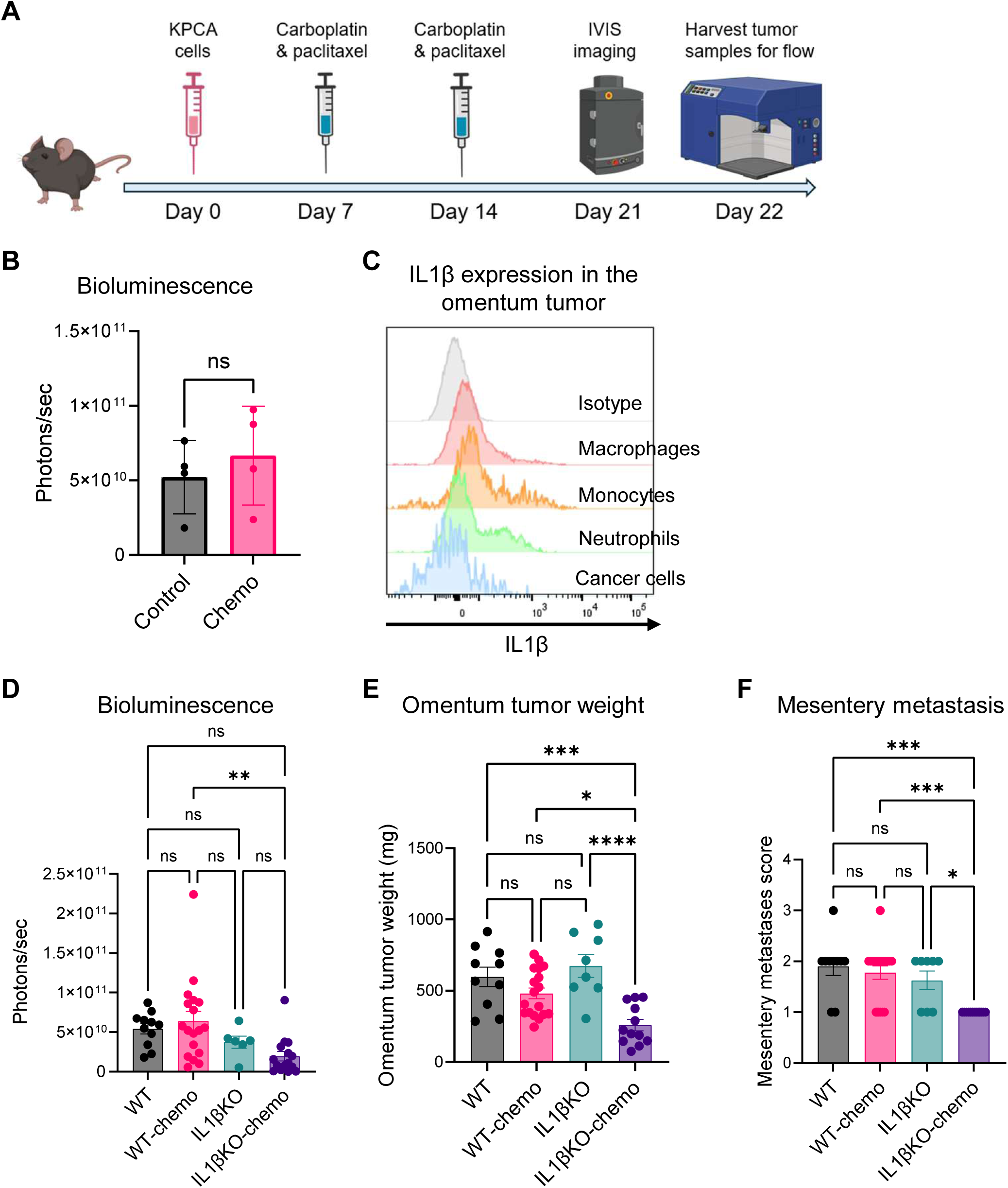
IL1β from myeloid cells contributes to OC chemoresistance (A) Experimental schematic to illustrate the timeline in KPCA model. (B) Quantification of bioluminescent signal in wild type (WT) mice treated with chemotherapy and vehicle control. (C) Intracellular staining of IL1β in myeloid cells and cancer cells from the omentum tumor. (D-F) Quantification of (D) bioluminescent signal, (E) omentum tumor weight and (F) mesentery metastasis in WT and IL1β KO mice treated with chemotherapy and vehicle control. Data from two or more independent runs combined and presented in bar graphs. Each dot represents one mouse (n=6 or more in each group). Student’s *t* test and One-way ANOVA with Tukey multiple comparisons test were used in (B) and (D-F), respectively. *P < 0.05; **P < 0.01; ***P < 0.001; ****P < 0.0001. Error bars are standard errors of the mean.

To investigate the functional role of myeloid cell–derived IL1β in chemotherapy, we compared chemotherapy responses between wild-type (WT) and IL1β knockout (IL1β KO) mice. While WT mice showed minimal response to chemotherapy, IL1β KO mice exhibited trend toward reduced bioluminescence signals and significant reductions in omental tumor burden and mesenteric dissemination following treatment (Figure 2D–2F). Notably, in the absence of chemotherapy, tumor growth of KPCA cells did not differ between WT and IL1β KO mice. A consistent result for omental tumor burden was observed using another CCNE1-overexpressed, HR-proficient cell line, BPCA (Supplementary Figure S2A and S2B). These results collectively suggest that IL1β derived from myeloid cells contributes to chemotherapy resistance in OC.

### IL1β recruits neutrophil and impairs chemotherapy response

To determine whether IL1β affects chemotherapy sensitivity through a direct or indirect mechanism, we first examined the direct action of IL1β on tumor cells. *In vitro* drug sensitivity assays were performed in the presence or absence of IL1β. The results showed that the IC50 values for carboplatin and paclitaxel remained largely unchanged regardless of IL1β’s presence or concentration, indicating that IL1β does not directly alter the chemosensitivity of OC (Figure 3A and 3B). To investigate the indirect effects of IL1β on chemosensitivity, we examined differences in the TME with and without chemotherapy in wild-type (WT) and IL1β KO mice. Flow cytometric analysis of immune cell populations in KPCA-derived omental tumors revealed that neutrophil infiltration was increased following chemotherapy in WT mice, whereas this increase was absent in IL1β KO mice (Figure 3C). The proportions of CD8⁺ T cells, CD4⁺ T cells, monocytes, and macrophages remained unchanged (Figure 3D–3G). Similarly, in BPCA tumors, omental lesions from IL1β KO mice exhibited significantly fewer neutrophils after chemotherapy compared to those from WT mice (Supplementary Figure S2C–2G). These findings suggest that IL1β contributes to the chemotherapy-induced recruitment of neutrophils.

**Figure 3.**
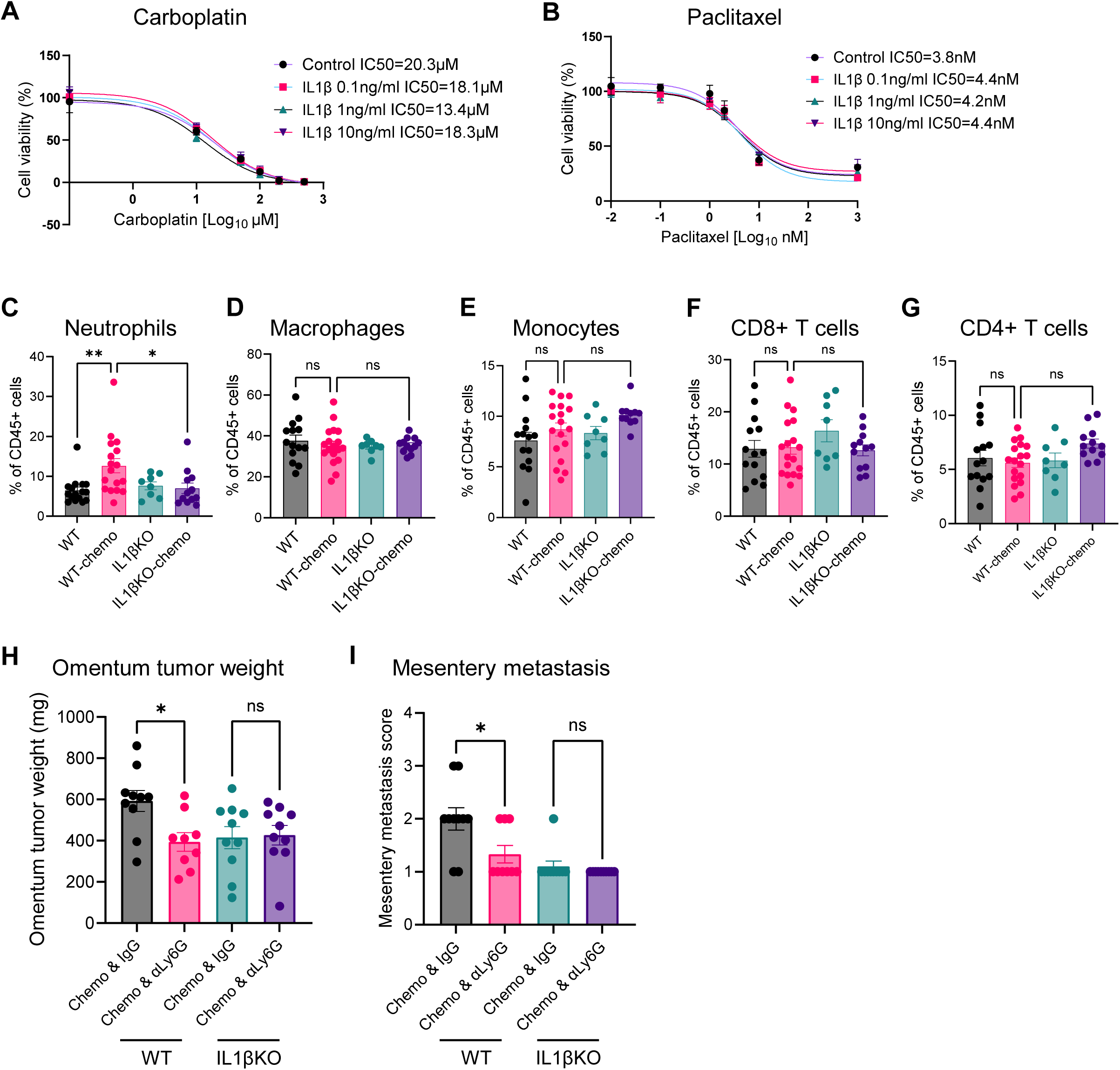
IL1β recruits neutrophil and impairs chemotherapy response (A and B) (A) Carboplatin and (B) paclitaxel IC_50_ assay of KPCA cells in the presence of IL1β at different concentrations. (C-G) Quantification of frequencies of (C) neutrophils, (D) macrophages, (E) monocytes, (F) CD8^+^ T cells, and (G) CD4^+^ T cells in omentum tumors treated as indicated and determined by flow cytometry. (H and I) (H) omentum tumor weight and (I) mesentery metastasis in WT and IL1β KO mice treated with chemotherapy together with αLy6G antibody and control IgG. In vitro data are representative of at least three independent experiments. In vivo data combined from three or more independent runs plotted where each dot represents one mouse (n=8 or more in each group). One-way ANOVA with Tukey multiple comparisons test were used in (C-I). *P < 0.05; **P < 0.01. Error bars are standard errors of the mean.

We next examined whether the IL1β–driven increase in neutrophils contributes to chemotherapy resistance. Indeed, neutrophil depletion under chemotherapy led to a reduction in omental tumor size and mesenteric dissemination compared to control-treated mice in WT mice. In contrast, no such differences were observed in IL1β KO mice (Figure 3H and 3I). These findings suggest that chemotherapy-induced neutrophil accumulation contributes to IL1β–mediated chemoresistance.

### T cell suppression and NETs contribute to the IL1β-neutrophil-induced chemoresistance

To investigate the mechanism by which chemotherapy-induced neutrophil accumulation contributes to chemoresistance, we performed single-cell RNA sequencing (scRNA-seq) of omental tumors from chemotherapy-treated WT and IL1β KO mice. Focusing on T cells and NK cells, the key components of antitumor immunity, we identified ten distinct clusters (Figure 4A, Supplementary Figure S3A). In IL1β KO mice, there was a marked reduction in cluster 0, which corresponds to stressed CD8 T cells (32) (Supplementary Figure S3B), and a dramatic enrichment of cluster 2, characterized by an effector CD8 phenotype expressing *Ifng*, *Tnf*, low levels of *Tox* (Figure 4B), and genes encoding proteins that reflect recent antigen experience, such as *Nr4a2*, *Nr4a3*, and *Tnfsf11* (33,34) (Supplementary Figure S3C). These findings suggest enhanced CD8 T cell–mediated antitumor immunity in the absence of IL1β. In contrast, comparative transcriptomic analysis of neutrophils between the two groups revealed less differences compared to those of T cells (Supplementary Figure S3D). Additionally, we confirmed that *Il1b* was only expressed in myeloid cells, i.e., neutrophils and macrophages in omentum tumors in our model (Supplementary Figure S3E).

**Figure 4.**
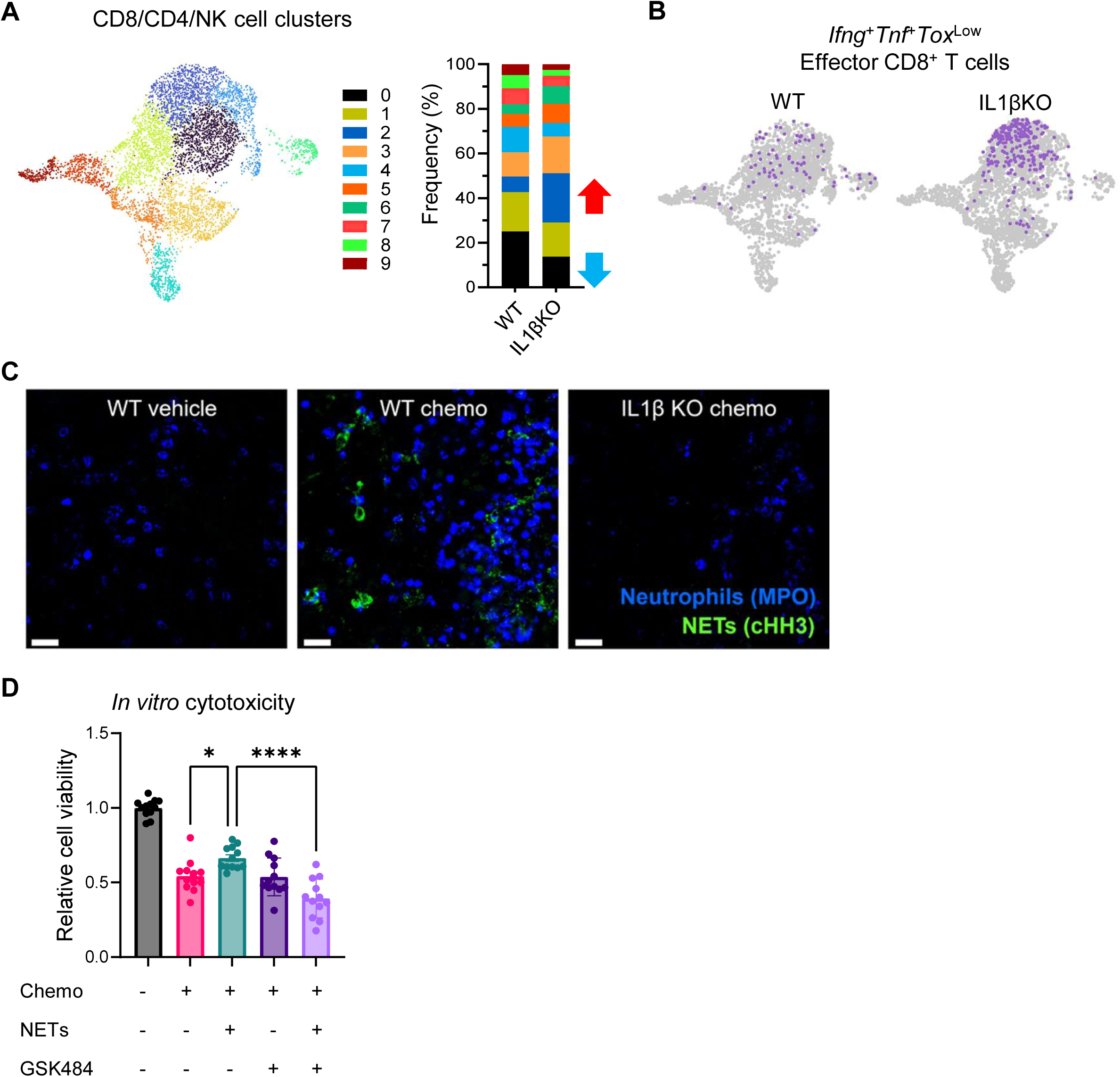
T cell suppression and NETs may contribute to the IL1β-neutrophil-induced chemoresistance (A) UMAP visualization of T/NK cells (k = 10 clusters) derived from chemotherapy-treated omental tumors of WT and IL1βKO mice, with the corresponding cluster frequency distribution. (B) UMAP visualization of the *IFNγ*^+^*TNF*^+^*Tox*^low^ effector CD8 T cell population in WT and IL1β KO mice. (C) representative confocal images of omentum tumors from WT and IL1β KO mice treated with chemotherapy and vehicle control. Positive cells were stained blue (MPO; neutrophil) and green (cHH3; NETs). (D) Cell viability assay with chemotherapeutic agents in NETs-conditioned media. In some conditions, GSK484 (NETs inhibitor; SML1658, Sigma) was added during NETs stimulation. Data combined from three independent runs (n=12 in each group). *P < 0.05; ****P < 0.0001. Error bars are standard errors of the mean.

We next investigated the role of neutrophil extracellular traps (NETs). NETs are extracellular DNA–protein networks released during a form of neutrophil cell death and have been reported to be induced following chemotherapy and to promote tumor progression in breast cancer (31). As NETs cannot be identified through scRNA-seq, we evaluated their presence by imaging of omentum tumor sections. In WT mice, chemotherapy induced both neutrophil infiltration and prominent NETs formation. These effects were abrogated in IL1β KO mice, indicating that IL1β is required for chemotherapy-induced NETs induction (Figure 4C). To determine whether NETs influence chemotherapy sensitivity, we conducted *in vitro* assays. The cytotoxicity of carboplatin and paclitaxel were attenuated in the presence of NETs, whereas pharmacologic inhibition of NETs reversed this resistance (Figure 4E). These findings suggest that IL1β– driven neutrophils contribute to chemoresistance during chemotherapy through both suppression of T cell– mediated antitumor immunity and promotion of NETs formation.

### IL1β induces neutrophil recruitment via CXCL2 upregulation after chemotherapy

We next investigated the mechanism by which IL1β promotes neutrophil recruitment following chemotherapy in OC tissues. Among neutrophil-related chemokines and growth factors, comparison of pre-and post-chemotherapy samples across publicly available datasets revealed that *CXCL2* was consistently upregulated after treatment in all datasets examined (Figure 5A). To identify the cellular targets of IL1β, we analyzed the expression of its receptor, IL1R1, using the same human post-chemotherapy scRNA-seq dataset as previously described (Figure 1G). *IL1R1* was found to be highly expressed in mesenchymal cell populations, particularly fibroblasts and mesothelial cells (Supplementary Figure S3F). *CXCL2* was also found to be highly expressed in both mesenchymal and macrophage populations (Supplementary Figure S3G). Interestingly, co-expression of *IL1R1* and *CXCL2* was mainly observed in fibroblast populations (Figure 5B). High IL1R1 expression in mesenchymal cells –particularly fibroblasts– was further confirmed by flow cytometry of omental tumors from the murine KPCA model (Figure 5C). Notably, in IL1β KO mice, *Cxcl2* expression in omental fibroblasts was significantly reduced following chemotherapy compared to WT controls (Figure 5D). TIMER analysis of the TCGA-OV cohort revealed that higher expressions of *IL1B*, *IL1R1*, and *CXCL2* significantly correlate with increased estimated neutrophil infiltration (Figure 5E). These results suggest that IL1β acts on IL1R1-expressing mesenchymal cells to induce CXCL2, thereby promoting neutrophil recruitment in response to chemotherapy.

**Figure 5.**
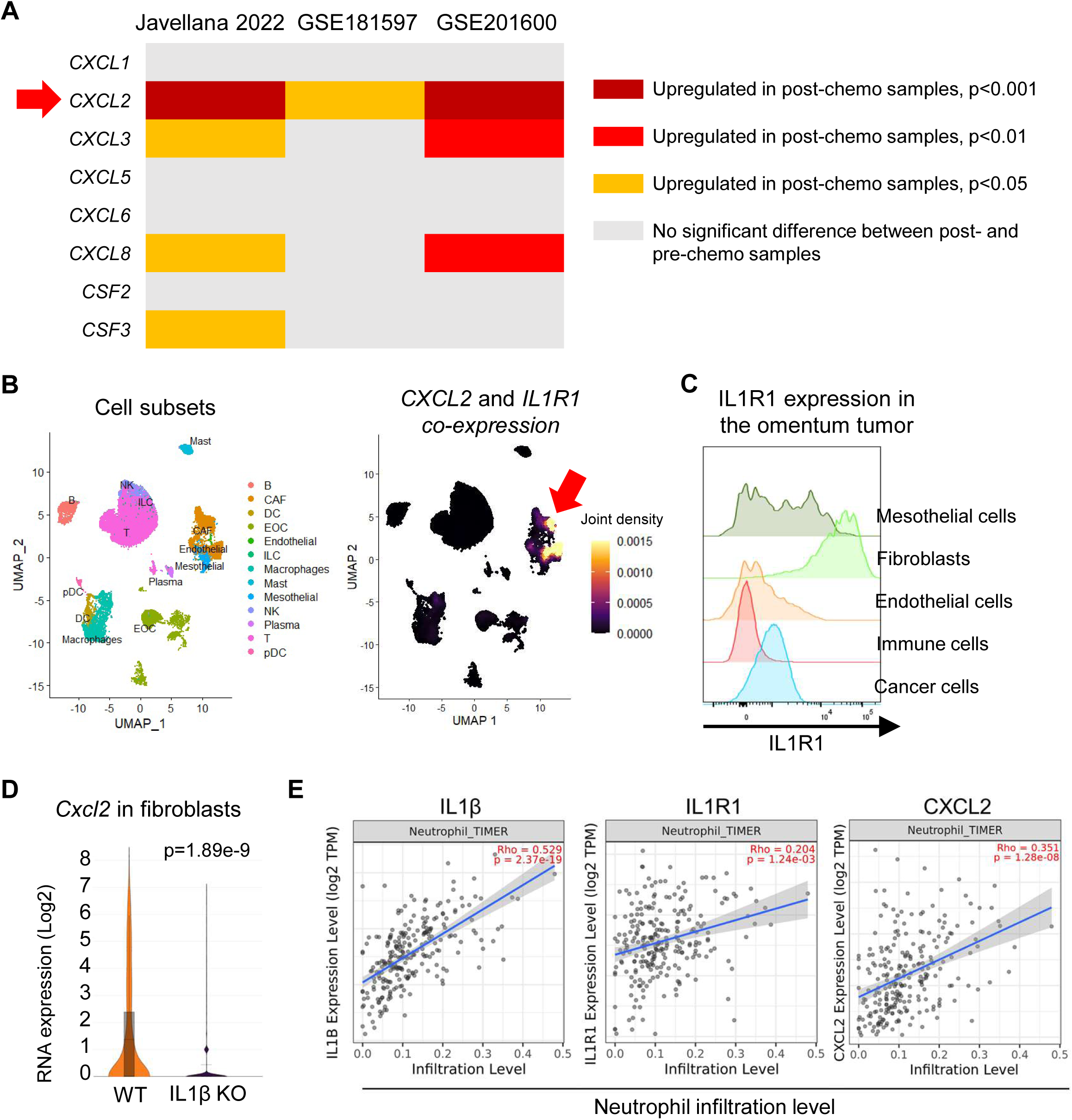
IL1β induces neutrophil recruitment via CXCL2 upregulation after chemotherapy (A) Comparison in cytokine expression related to neutrophil migration and differentiation between pre-and post-chemotherapy samples using public datasets. (B) UMAP plot of cells from human OC post-chemotherapy tumor samples (GSE165897) colored by cell subsets (left) and *IL1R1* and *CXCL2* co-expression (right). (C) IL1R1 staining in mesenchymal cells, immune cells and cancer cells from the omentum tumor. (D) Violin plot showing *Cxcl2* expression in fibroblasts in the omentum tumor from WT and IL1β KO mice from the mouse scRNAseq data. (E) Correlation between estimated neutrophil infiltration and *IL1B*, *IL1R1* and *CXCL2* expression in human OC tumor samples using TIMER.

### IL1R1 KO mice exhibit chemo-sensitivity

As IL1R1 is predominantly expressed in host-derived mesenchymal cells and IL1β does not have any impact on chemoresistance in OC cells in vitro, we next investigated whether host IL1R1 knockout (IL1R1 KO) could phenocopy IL1β KO by abrogating chemotherapy-induced neutrophil recruitment and the associated chemoresistance. Indeed, IL1R1 KO mice exhibited reduced omental tumor burden and decreased mesenteric metastasis following chemotherapy compared to WT controls (Figure 6A–6C). Consistent with these findings, the number of neutrophils was lower in IL1R1 KO mice after chemotherapy than in WT mice, whereas the proportions of other immune cell populations remained unchanged (Figure 6D–6H). Moreover, functional analysis of T cells revealed enhanced IFNγ and TNF production in CD8⁺ T cells isolated from the omentum of IL1R1 KO mice (Supplementary Figure S4A and S4B). Collectively, these results support the involvement of the IL1β–IL1R1–neutrophil axis in promoting chemotherapy resistance in OC.

**Figure 6.**
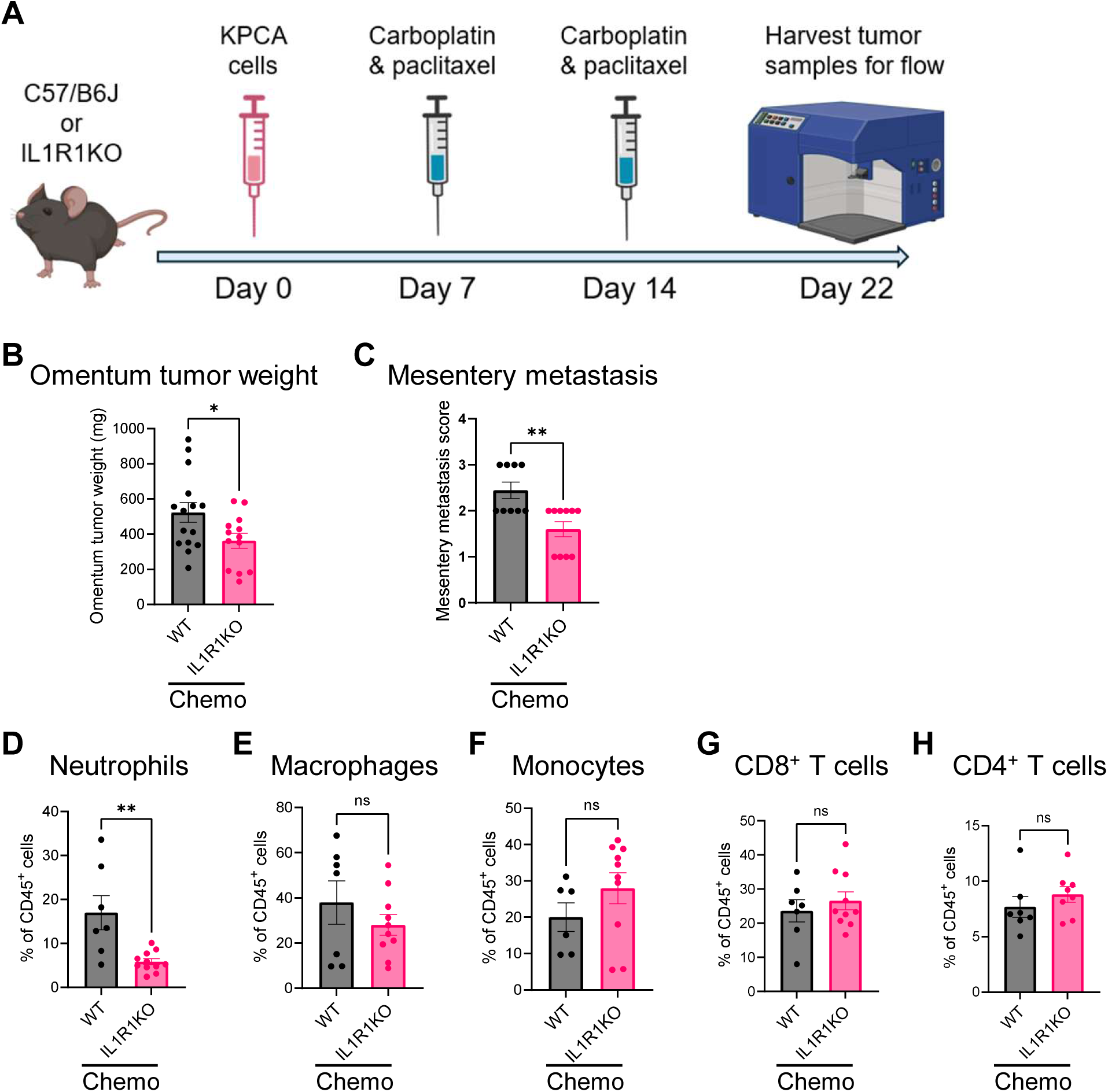
IL1R1 KO mice exhibit chemo-sensitivity (A) Experimental schematic to illustrate the timeline in KPCA model in WT and IL1R1 KO mice. (B-C) Quantification of (B) omentum tumor weight and (C) mesentery metastasis in WT and IL1R1 KO mice treated with chemotherapy. (D-H) Quantification of frequencies of (D) neutrophils, (E) macrophages, (F) monocytes, (G) CD8^+^ T cells, and (H) CD4^+^ T cells in omentum tumors treated as indicated and determined by flow cytometry. (B-H) In vivo data combined from two or more independent runs plotted where each dot represents one mouse (n=7 or more in each group). *P < 0.05; **P < 0.01. Error bars are standard errors of the mean.

### Neutrophils and NETs are increased in human post-chemo OC samples

Finally, we compared neutrophils and NETs in human OC tissues before and after chemotherapy. Paired omental samples were analyzed from 16 patients who underwent sampling of the omentum for diagnostic purposes and subsequently underwent interval debulking surgery including omentectomy, in which residual tumors were present in the omentum. Patient characteristics are summarized in Supplementary Table S2. All patients received neoadjuvant chemotherapy consisting of carboplatin and taxane, with a median of three treatment cycles. The median interval between the last chemotherapy dose and interval debulking surgery was 28 days. To quantify each cellular population within the tumors, we employed the pathology AI platform HoVer-NeXt for cell-type annotation on H&E-stained slides (Figure 7A, Supplementary Figure S5A and S5B). The region of interest (ROI) included the entire slide area. At higher magnification, cells labeled as neutrophils appeared smaller than tumor cells and exhibited segmented nuclei (Figure 7B). The proportion of cancer cells among total cells did not change before and after chemotherapy (Figure 7C). In contrast, the proportion of neutrophils among total cells and among tumor cells increased after chemotherapy (Figure 7D and 7E). Notably, circulating neutrophil counts decreased following chemotherapy (Supplementary Figure S5C). Next, we evaluated NETs using multiplex immunofluorescence staining for cHH3 and MPO (Figure 7F). A trend toward an increased proportion of NETs among total cells was observed following chemotherapy (Figure 7G). Collectively, these findings suggest that chemotherapy induces infiltration of neutrophils and formation of NETs in human OC.

**Figure 7.**
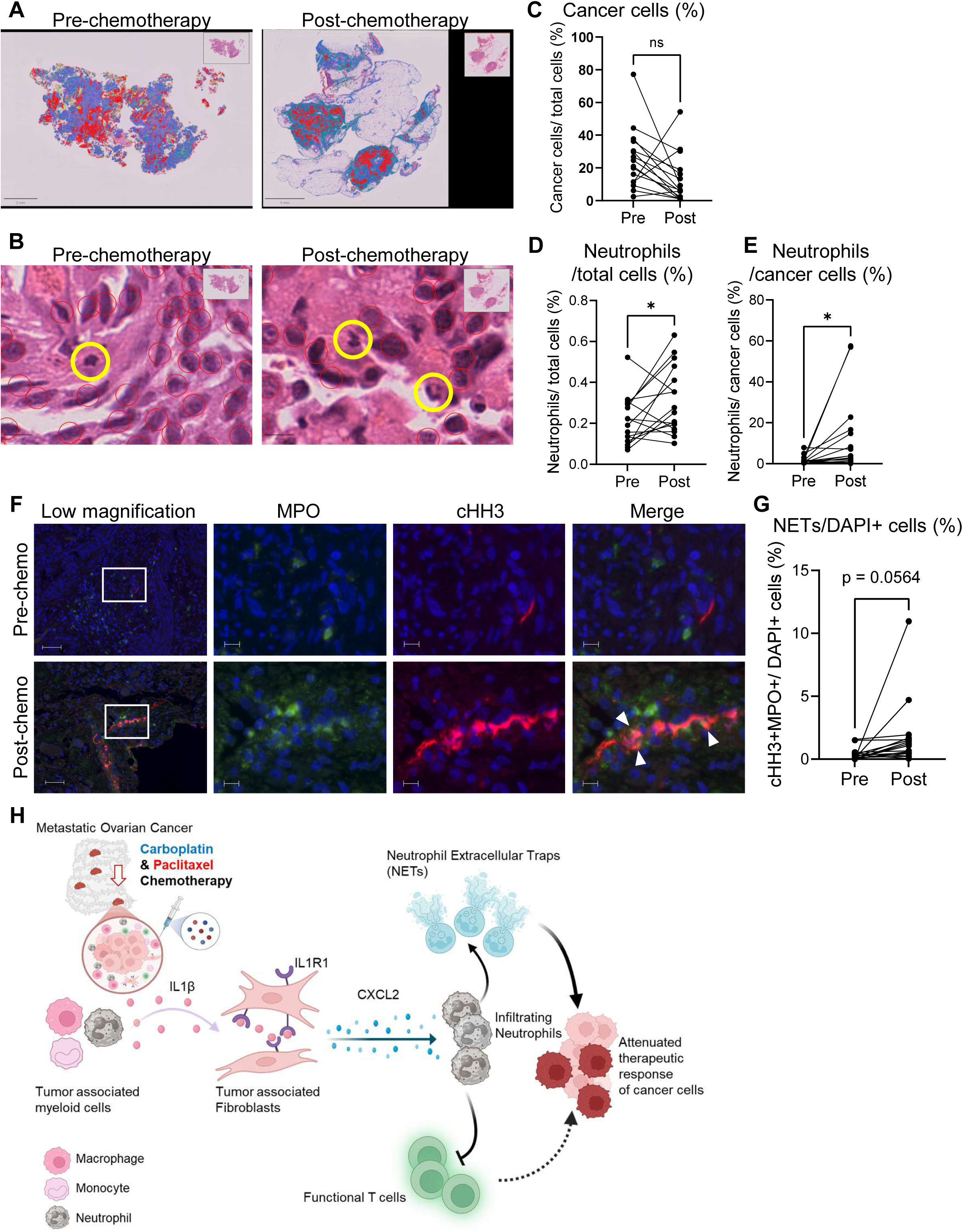
Neutrophils and NETs are increased in human post-chemotherapy OC samples (A) HoVer-NeXt AI model analyzed whole-slide images of pre-and post-chemotherapy specimens from same patient. Cells are color-coded as follows: red, cancer cells; blue, connective tissue cells; yellow, neutrophils; green, lymphocytes; orange, eosinophils; and teal, plasma cells. Scale bars: 2 mm (pre-chemotherapy) and 5 mm (post-chemotherapy). (B) Higher-magnification images of (A). Neutrophils (yellow circles) and cancer cells (red circles) are indicated. Scale bars, 10 µm. (C) Proportion of cancer cells within the total cell population in pre-and post-chemotherapy specimens. (D, E) Neutrophil frequency as a percentage of (D) total cells and (E) cancer cells in pre-and post-chemotherapy specimens. (F) Representative immunofluorescence images of pre-and post-chemotherapy specimens. Positive cells were stained green (MPO), red (cHH3) and blue (DAPI). Scale bars: 50 μm (lower-magnification) and 10 μm (higher-magnification). (G) NETs frequency as a percentage of DAPI+ cells in pre-and post-chemotherapy specimens. (H) Graphical abstract. The figure is created using BioRender and Adobe illustrator. Paired *t* test was used in (C-E and G). *P < 0.05.

## Discussion

We identified chemotherapy-induced inflammatory remodeling of the tumor microenvironment— specifically the role of myeloid cell–derived IL1β—as a contributor to chemoresistance in HGSC. Following chemotherapy, IL1β promotes neutrophil recruitment through CXCL2 induction in mesenchymal cells – particularly fibroblasts–, leading to the formation of NETs and suppression of T cell–mediated antitumor immunity, thereby facilitating chemoresistance (Figure 7H). These phenotypes were markedly attenuated in mice deficient in either IL1β or its receptor IL1R1, resulting in improved sensitivity to chemotherapy. To our knowledge, this study is the first to demonstrate that therapy-induced inflammation can extrinsically drive chemoresistance in OC.

IL1β is a proinflammatory cytokine known to promote tumor initiation and progression. However, in the present study, we observed no difference in tumor growth between untreated IL1β KO and WT mice. IL1β contributes to *in vivo* tumor progression only in the context of chemotherapy in our models (CCNE1 and Akt2 overexpressed) (Figure 2D–2F, Supplementary Figure S2B). This finding contrasts with observations in other malignancies such as breast cancer (35), pancreatic cancer (36), and glioblastoma (37), where host IL1β knockout or IL1β blockade alone has been sufficient to suppress tumor growth. These cancer type–specific discrepancies indicate that the function of IL1β is context-and tissue-dependent.

In order to determine whether IL1β directly modulates chemotherapy response, we tested OC cells *in vitro* and found no significant change in their sensitivity in the presence of IL1β (Figure 3A and 3B). This differs from previous studies in other cancers, which showed that IL1β inhibits etoposide-induced apoptosis in pancreatic cancer (38) and promotes carboplatin resistance via ZEB1 activation in colorectal cancer (39). In contrast, our study demonstrates that IL1β promotes chemoresistance indirectly, through host IL1β–IL1R1 signaling that induces neutrophil recruitment (Figure 3H, 3I and 5A–5D). A similar mechanism has been described in breast cancer, where chemotherapy-induced IL1β expression contributes to resistance via neutrophil and NETs induction (31), highlighting the significance of IL1β– mediated indirect effects on treatment sensitivity.

Among the mechanisms by which neutrophils promote tumor progression, suppression of T cell function has been well documented in OC and other malignancies (40–42). In the present study, tumors from IL1β KO or IL1R1 KO mice failed to exhibit chemotherapy-induced neutrophil recruitment and instead showed enhanced T cell–mediated immune activation compared to tumors from WT mice (Figure 4A and 4B and Supplementary Figure S3 and S4). These findings suggest that neutrophil-mediated suppression of T cell immunity under chemotherapy may contribute to IL1β-mediated treatment resistance. Interestingly, we observed that chemotherapy enhances neutrophil infiltration and NETs formation in both human and murine HGSC tumors, and that NETs contribute to chemoresistance (Figure 4C and 4D, Figure 7). Chemoresistance mediated by NETs has been attributed to multiple mechanisms, including the stimulation of TGF-β signaling that drives Epithelial-mesenchymal transition (EMT) (31), and the internalization of NETs-derived DNA by tumor cells, which impairs drug–DNA interactions within the nucleus (43).

In OC, immune checkpoint inhibitors have shown limited clinical efficacy, and their combination with chemotherapy has also failed to demonstrate significant therapeutic benefit (44,45). Consequently, no immunological strategies have yet been established to overcome treatment resistance in OC. The lack of additional benefit from the anti–IL1β antibody canakinumab in combination with chemotherapy in lung cancer is disappointing (46,47). However, the absence of *IL1B* upregulation following chemotherapy in lung cancer (48), along with the tissue-specific nature of IL1β activity, does not preclude the potential efficacy of IL1β inhibition in OC. Furthermore, with regard to NETs inhibition, a first-in-human phase I clinical trial of the humanized monoclonal antibody CIT-013—designed to target citrullinated histones H2A and H4—has been reported for the treatment of autoimmune diseases (49). These novel agents targeting NETs may offer a promising therapeutic strategy to overcome chemotherapy resistance, warranting further investigation.

The limitation of this study is that the murine models used do not fully capture the intratumoral heterogeneity of human HGSC, which is genetically unstable and characterized by the presence of subclones harboring distinct genotypes (50). The mouse models employed in this study were generated using clonal cell lines with uniform genetic alterations and therefore do not fully recapitulate the heterogenous TME of human HGSC. Although bulk RNA sequencing of human HGSC samples demonstrated upregulation of *IL1B* following chemotherapy, this study does not address how *IL1B* expression, neutrophil infiltration, or neutrophil function may vary across spatially distinct subclones within the tumor. Future studies employing spatial transcriptomics or genomics will be essential to delineate how intratumoral heterogeneity and local immune landscapes evolve in response to chemotherapy in HGSC.

In conclusion, we demonstrate that chemotherapy-induced IL1β–dependent neutrophil infiltration contributes to chemoresistance in OC, uncovering a paradoxical mechanism by which therapy-induced inflammation undermines therapeutic efficacy. Targeting the IL1β–neutrophil–NETs axis may represent a promising strategy to overcome chemoresistance in OC.

## Author’s contributions

T. Miyamoto: Conceptualization, Data curation, Formal analysis, Funding acquisition, Investigation, Methodology, Project administration, Visualization, Writing - original draft, Writing - review & editing, Y. Ye: Investigation, Methodology, Validation, Writing - original draft, Writing - review & editing, B.S. Manning: Investigation, Methodology, B. Murphy: Formal analysis, Investigation, Visualization, M. Minakuchi: Investigation, Methodology, K. Hamada: Data curation, Formal analysis, Writing - review & editing, R. Mizuno: Resources, Writing - review & editing, M. Taki: Resources, Writing - review & editing, K. Yamanoi: Resources, Writing - review & editing, R. Murakami: Resources, Writing - review & editing, M. Mandai: Resources, Writing - review & editing, Y. Ye: Data curation, Formal analysis, Writing - review & editing, J. Wickramasinghe: Data curation, Formal analysis, Writing - review & editing, Y. Nefedova: Resources, A. Kossenkov: Data curation, Formal analysis, Software, Validation, Visualization, Writing - review & editing, N. Zhang: Conceptualization, Data curation, Formal analysis, Funding acquisition, Investigation, Methodology, Project administration, Supervision, Validation, Visualization, Writing - original draft, Writing - review & editing.

## Acknowledgements

We thank Wistar core facilities (imaging, histology, flow cytometry, animal facility, and genomics). This study was supported by Concern Foundation (N. Zhang), V Foundation (V2024-026; N. Zhang), Department of Defense Ovarian Cancer Research Program (HT94252410206; N. Zhang), and Japan Society for the Promotion of Science (25K20087; T. Miyamoto).

---

**Supplementary Figure S1.**
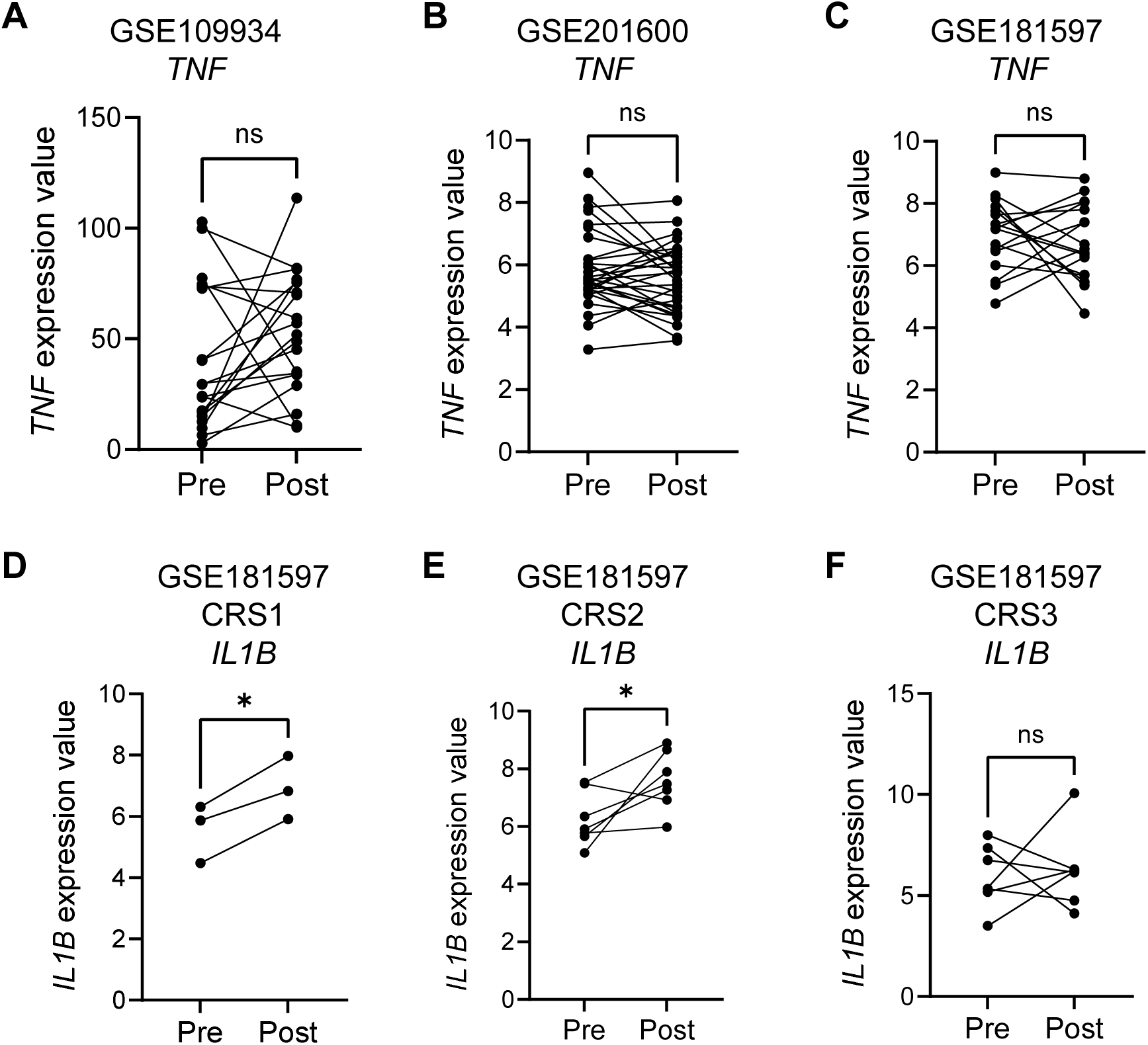
IL1β is upregulated in human OC after chemotherapy, related to Figure 1 (A-C) Comparison of *TNF* gene expression between pre-and post-chemotherapy samples using (A) GSE109934, (B) GSE201600 and (C) GSE181597 datasets. (C-F) Comparison of *IL1B* gene expression between pre-and post-chemotherapy samples in the GSE181597 dataset, stratified by chemotherapy response score (CRS). Paired *t* test was used. *P < 0.05.

**Supplementary Figure S2.**
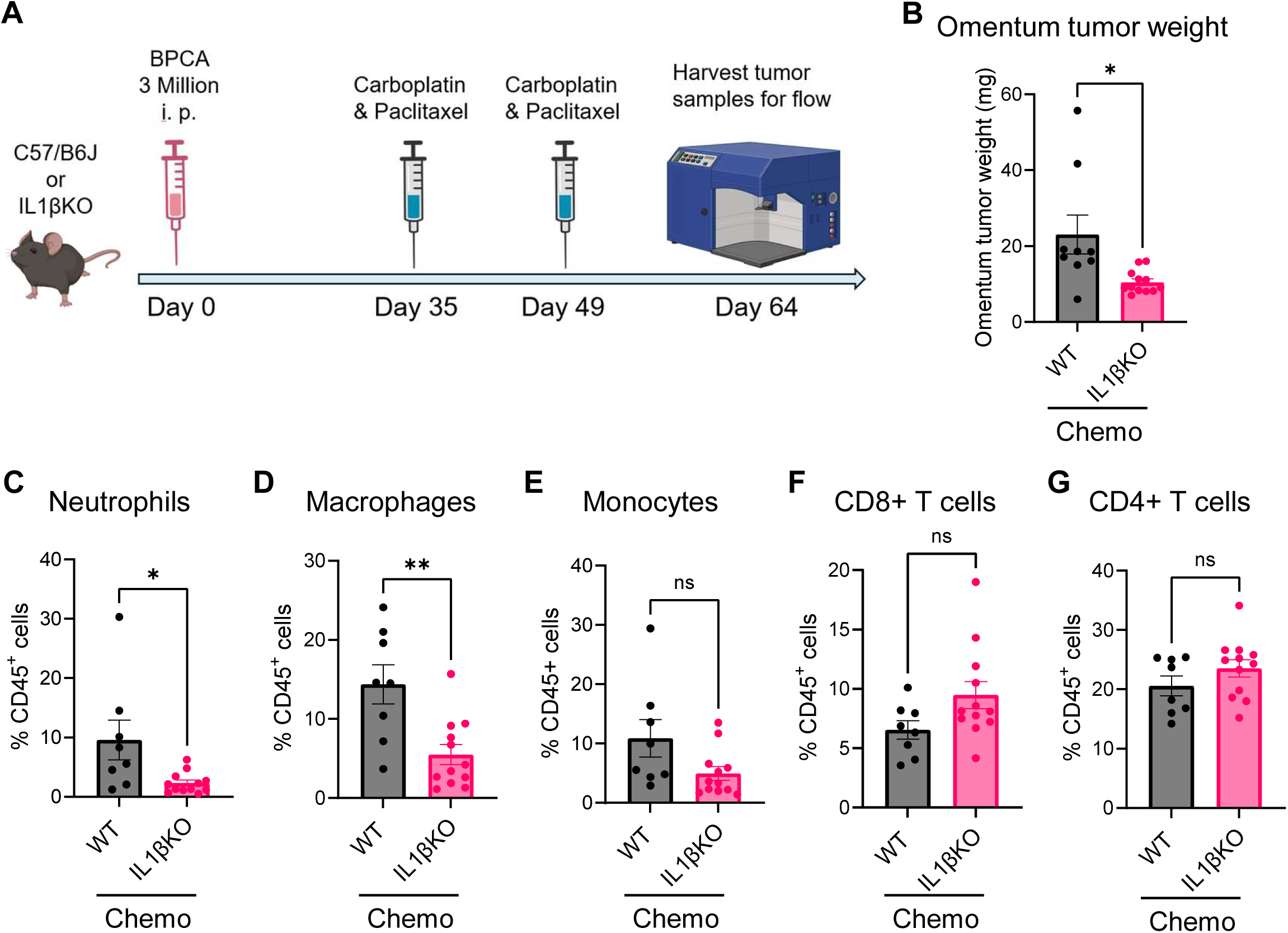
BPCA-chemo model in IL1bKO mice, related to Figure 2 and 3 (A) Experimental schematic to illustrate the timeline in BPCA model in WT and IL1β KO mice. (B) Quantification of omentum tumor weight in WT and IL1β KO mice treated with chemotherapy. (C-G) Quantification of frequencies of (C) neutrophils, (D) macrophages, (E) monocytes, (F) CD8^+^ T cells, and (G) CD4^+^ T cells in omentum tumors determined by flow cytometry. In vivo data combined from two or more independent runs plotted where each dot represents one mouse (n=7 or more in each group). *P < 0.05; **P < 0.01. Error bars are standard errors of the mean.

**Supplementary Figure S3.**
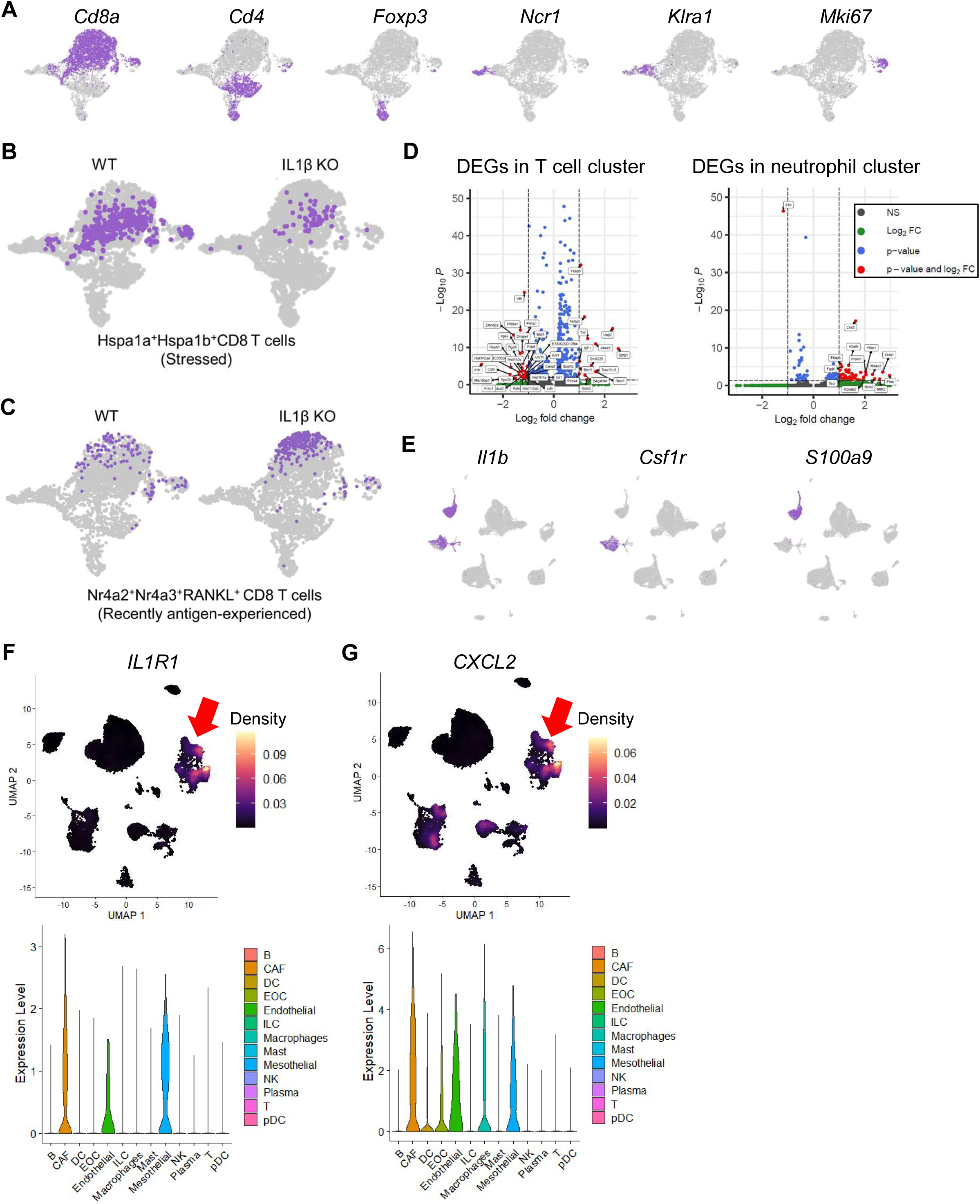
scRNAseq analysis of murine and human OC samples, related to Figure 4 and 5 (A) UMAP projection of T/NK cell clusters highlighting cells expressing canonical subset markers. (B and C) UMAP projection of T/NK cell clusters highlighting (B) stressed and (C) recently antigen-experienced CD8 T cells derived from chemotherapy-treated omental tumors of WT and IL1βKO mice. (D) Volcano plots depicting differential gene expression between WT and IL1β KO mice within T cell (left) and neutrophil (right) populations. (E) UMAP projection of immune cell clusters highlighting expression of *Il1b*, *Csf1r*, and *S100a9*. (F) Density plots (top) illustrating *IL1R1*-expressing populations and violin plots (bottom) showing subset-specific *IL1R1* expression in post-chemotherapy tumors. (G) Density plots (top) illustrating *CXCL2*-expressing populations and violin plots (bottom) showing subset-specific *CXCL2* expression in post-chemotherapy tumors.

**Supplementary Figure S4.**
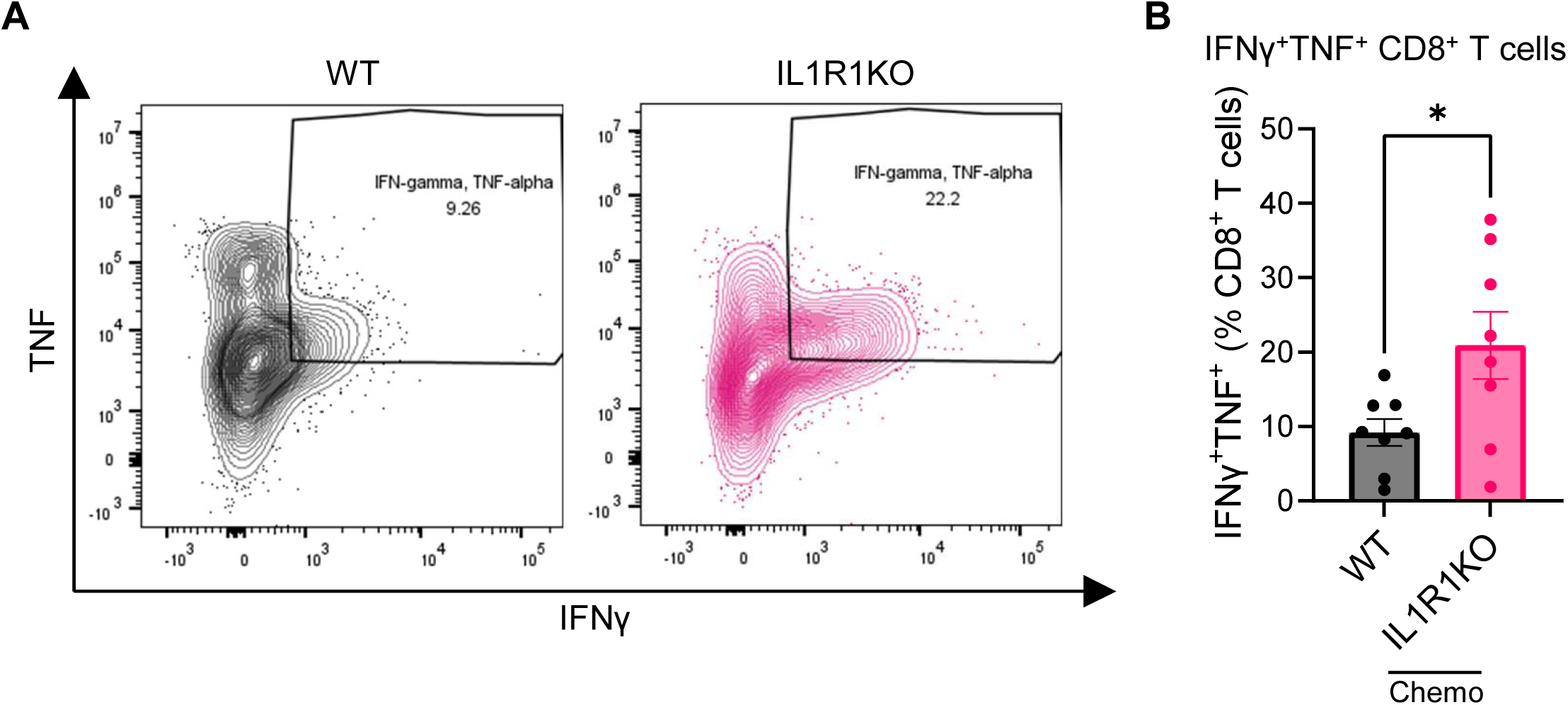
Enhanced IFNγ/TNF production in CD8 T cells isolated from the omentum of IL1R1 KO mice, related to Figure 6 (A and B) IFNγ/TNF in omental tumor CD8 T cells from WT and IL1R1 KO mice: (A) representative plots; (B) frequencies of IFNγ⁺/TNF⁺ cells. Data combined from two independent runs plotted where each dot represents one mouse (n=8 in each group). *P < 0.05. Error bars are standard errors of the mean.

**Supplementary Figure S5.**
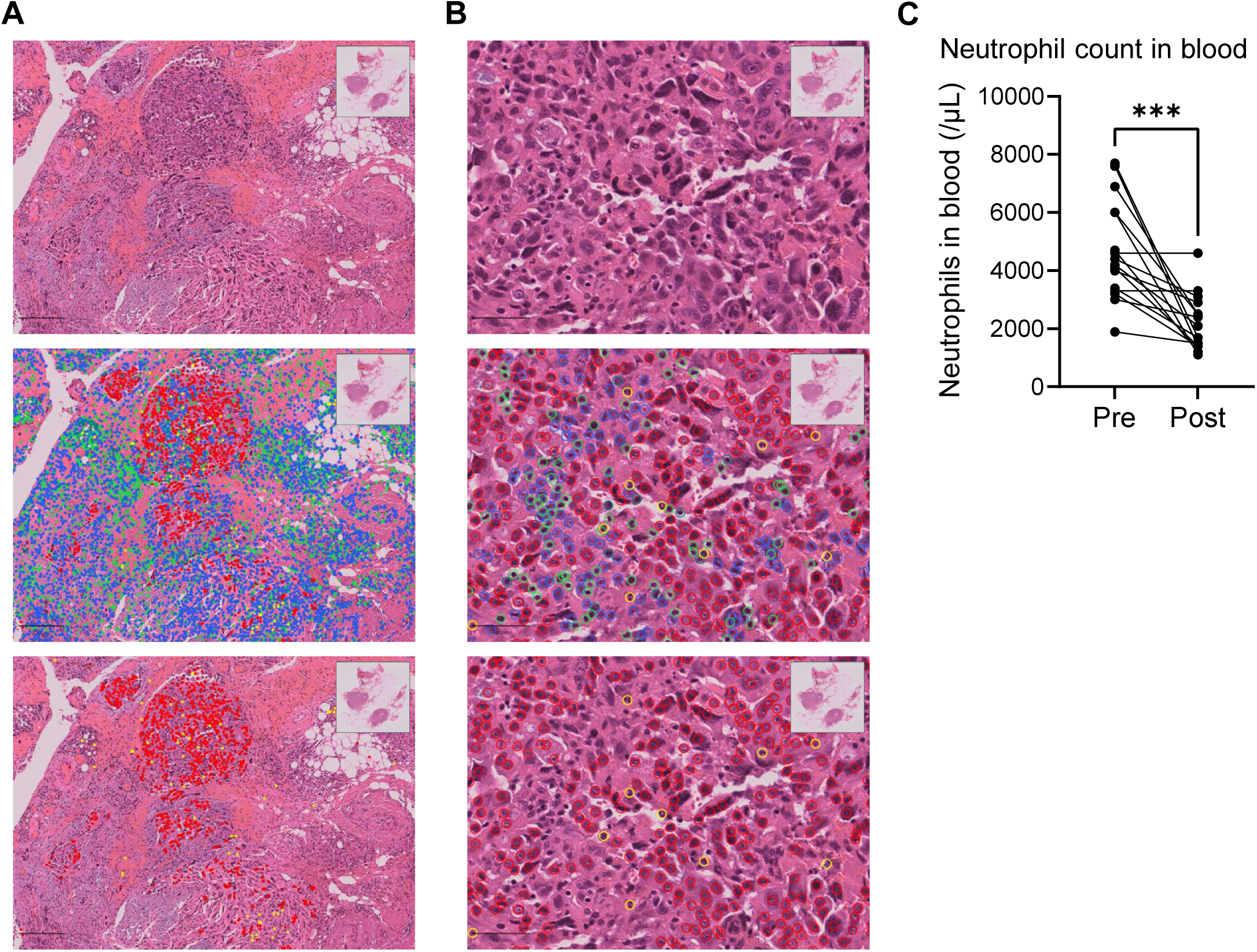
Changes in neutrophil between pre-and post-chemotherapy human OC samples, related to Figure 7 (A and B) HoVer-NeXt analyzed H&E slides at (A) lower and (B) higher magnification. Top: original, unannotated images. Middle: cell-type overlays (red, cancer cells; blue, stromal cells; yellow, neutrophils; green, lymphocytes; orange, eosinophils; teal, plasma cells). Bottom: annotations highlighting only neutrophils (yellow) and cancer cells (red). Scale bars: 200 μm (A) and 50 μm (B). (C) Paired comparison of peripheral-blood absolute neutrophil counts measured before and after chemotherapy. Paired *t* test was used. ***P < 0.001.

